# Dihydroxyacetone phosphate signals glucose availability to mTORC1

**DOI:** 10.1101/2020.06.21.161927

**Authors:** Jose M. Orozco, Patrycja A. Krawczyk, Sonia M. Scaria, Andrew L. Cangelosi, Sze Ham Chan, Tenzin Kunchok, Caroline A. Lewis, David M. Sabatini

**Affiliations:** Whitehead Institute for Biomedical Research and Massachusetts Institute of Technology, Department of Biology, 455 Main Street, Cambridge, Massachusetts 02142, USA; Howard Hughes Medical Institute, Department of Biology, Massachusetts Institute of Technology, Cambridge, Massachusetts 02139, USA; Koch Institute for Integrative Cancer Research and Massachusetts Institute of Technology, Department of Biology, 77 Massachusetts Avenue, Cambridge, Massachusetts 02139, USA; Broad Institute of Harvard and Massachusetts Institute of Technology, 415 Main Street, Cambridge, Massachusetts 02142, USA

## Abstract

In response to nutrients, the mTOR complex 1 (mTORC1) kinase regulates cell growth by setting the balance between anabolic and catabolic processes. To be active, mTORC1 requires the environmental presence of amino acids and glucose, which provide the building blocks for the biosynthesis of most macromolecules. While a mechanistic understanding of amino acid sensing by mTORC1 is emerging, how glucose activates mTORC1 remains mysterious. Here, we used metabolically engineered human cells to identify glucose-derived metabolites required to activate mTORC1. We find that mTORC1 senses a metabolite downstream of the aldolase and upstream of the glyceraldehyde 3-phosphate dehydrogenase steps of glycolysis and pinpoint dihydroxyacetone phosphate (DHAP) as the key molecule. In cells expressing a triose kinase, the synthesis of DHAP from dihydroxyacetone is sufficient to activate mTORC1 even in the absence of glucose. Genetic perturbations in the GATOR-Rag signaling axis abrogate glucose sensing by mTORC1. DHAP is the glycolytic metabolite along with glyceraldehyde 3-phosphate (GAP) that has the greatest fold-change between cells in high and low glucose. DHAP is a precursor for lipid synthesis, a process under the control of mTORC1, which provides a potential rationale for the sensing of DHAP by mTORC1.

---

How organisms sense and adapt to the availability of nutrients in the environment is incompletely understood. One key pathway for doing so is the signaling system anchored by the mTOR Complex 1 (mTORC1) kinase, which regulates cell growth and metabolism in response to nutrients, such as amino acids and glucose^1^. Aberrant mTORC1 signaling is implicated in many diseases, including several associated with nutrient overload, such as diabetes and non-alcoholic steatohepatitis (NASH)^2–5^. In contrast to the mechanistic understanding beginning to emerge for how mTORC1 senses amino acids, much less is known about how it detects glucose.

Glucose deprivation can inhibit mTORC1 through a pathway that requires activation of the energy sensing kinase AMPK^6–8^, caused by a rise in the AMP to ATP ratio. However, there is also significant evidence that mTORC1 can sense glucose through an AMPK-independent route^9–12^. While potential models for this route have been proposed, several involving glycolytic intermediates and their cognate enzymes, many are mutually incompatible, and no consensus has emerged on either the mechanisms involved or the molecule, if not glucose, that is sensed^13–19^. Here, we metabolically engineered cells to probe the role of glucose-derived metabolites in the regulation of mTORC1. We find that a metabolite previously not connected to mTORC1 plays a key role in its activation by glucose.

## Results

To examine in our cell system the role of AMPK in glucose-sensing by mTORC1, we generated AMPKα1 and AMPKα2 double knockout (DKO) HEK-293T cells. As expected, in these cells glucose starvation did not increase the phosphorylation of the canonical AMPK substrate Acetyl CoA Carboxylase (ACC)^20^. Despite the absence of AMPK activity, glucose starvation still inhibited the phosphorylation of the mTORC1 substrate S6K1, albeit to a lesser degree than in wild-type cells (Fig 1A). These results support the existing notion that mTORC1 can sense glucose through at least two pathways, one AMPK-dependent and the other not. In order to focus on the AMPK-independent mechanism, we undertook all further experiments in the AMPK DKO cells.

**Figure 1.**
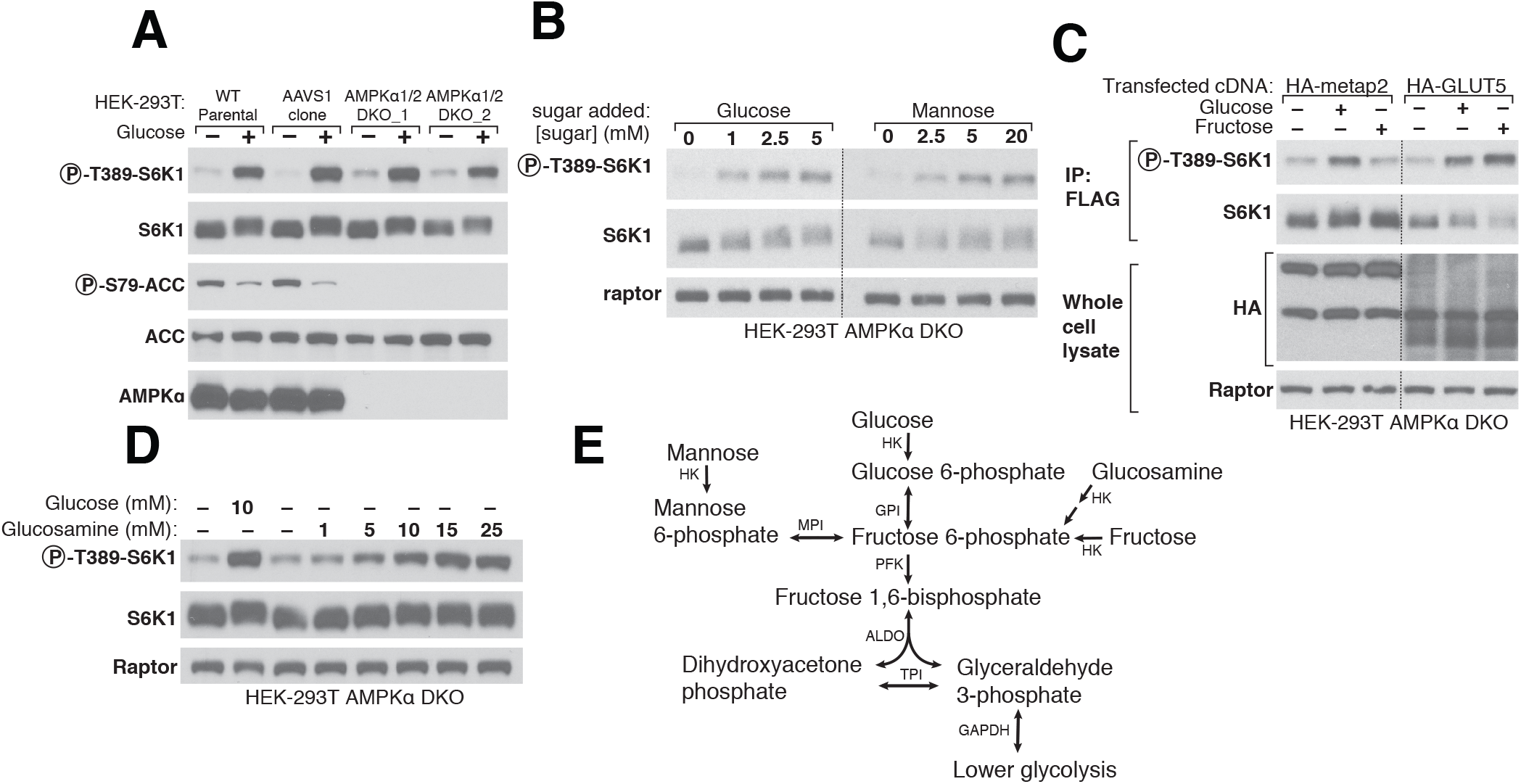
Various sugars can activate mTORC1 independently of AMPK. **A)** AMPK is not necessary for glucose to regulate mTORC1. Wildtype parental HEK-293T cells, a control cell line generated by expressing Cas9 and an sgRNA targeting the AAVS1 locus (AAVS1 clone), or two AMPKαdouble knockout (DKO) cell lines generated with sgRNAs targeting both genes encoding the AMPKαsubunit of AMPK (*PRKAA1 and PRKAA2*), were starved of glucose for 1 hour or starved and then re-stimulated with glucose for 10 minutes. Whole cell lysates were prepared and analyzed by immunoblotting using the indicated antibodies. **B)** In the absence of glucose, mannose activates mTORC1 in AMPKαDKO HEK-293T cells. Cells were incubated in media containing the indicated concentrations of glucose or mannose for 1 hour prior to the preparation and analysis of cell lysates. **C)** Fructose, in the absence of glucose, activates mTORC1 in cells expressing the fructose transporter GLUT5. FLAG-immunoprecipitates were prepared from cells co-expressing FLAG-S6K and either a control protein (HA-metap2) or HA-GLUT5. Immunoprecipitates and cell lysates were analyzed by immunoblotting for the phosphorylation state or levels of the indicated proteins. Dashed line indicates the splicing of two different exposures to show the relevant dynamic range of metap2 expressing cells or GLUT5 expressing cells. **D)** Glucosamine, in the absence of glucose, activates mTORC1 in AMPKαDKO HEK-293T cells. **E)** Schematic demonstrating how hexoses feed into glycolysis according to the KEGG pathway (HK: hexokinase; GPI: glucose 6-phosphate isomerase; PFK: phosphofructokinase; ALDO: Aldolase; TPI: triosephosphate isomerase; GAPDH: glyceraldehyde 3-phosphate dehydrogenase).

To probe the features of glucose necessary for it to activate mTORC1, we asked if glucose analogs as well as related sugars can substitute for glucose. While non-metabolizable (5-thio-glucose, 6-deoxyglucose, 2-deoxyglucose) and methylated (methyl α-D-glycopyranoside, 3-O-methyl-D-glucpyranose) glucose analogs could not (Fig S1A), the metabolizable sugars^21^ mannose, glucosamine, and fructose readily re-activated mTORC1 in glucose-starved cells (Fig. 1B-E). These results suggest that mTORC1 responds to a glucose-derived metabolite rather than to glucose itself, and that the pathway that generates the sensed molecule can be fed by fructose, mannose, or glucosamine. Given this, and that we found that the pentose phosphate pathway (PPP) is not required for glucose to activate mTORC1 (Fig S1B-C), we focused on the metabolism of glucose via glycolysis.

To pinpoint metabolites important for mTORC1 activation, we developed a genetic approach for inhibiting glycolytic enzymes because, other than for GAPDH^22,23^, there are few specific small molecule inhibitors targeting them. There are two key challenges to doing so: (1) glucose metabolism is essential in proliferating cells, and (2) many steps in glycolysis are catalyzed by enzymes with redundant paralogues. To overcome these issues, we built conditional knockout cell lines (‘dox-off’ cells) that lack all paralogues for a particular enzyme and express a complementing cDNA that is repressible by doxycycline.

Hexokinase phosphorylates glucose and mannose to generate glucose 6-phosphate (G6P) and mannose 6-phosphate (M6P), respectively, which specific isomerases (Glucose 6-phosphate isomerase (GPI) and mannose 6-phosphate isomerase (MPI)) convert through a reversible reaction into fructose 6-phosphate (F6P) (Fig 2A)^21^. In GPI dox-off cells, doxycycline treatment eliminated GPI expression and prevented glucose, but not mannose, from activating mTORC1 (Fig 2B). Metabolite profiling confirmed that glucose was not metabolized further than G6P in cells lacking GPI. In contrast, addition of mannose was able to restore the normal levels of all glycolytic metabolites except for G6P (Fig. 2A). Collectively, these data indicate that the glucose-derived metabolite that leads to mTORC1 activation must be made downstream of GPI, and rule out glucose itself and G6P as candidate signaling molecules. Our results differ from previous work in cardiomyocytes that proposed that G6P signals glucose levels to mTORC1^14,17–19^, suggesting that glucose sensing may differ between cell types.

**Figure 2.**
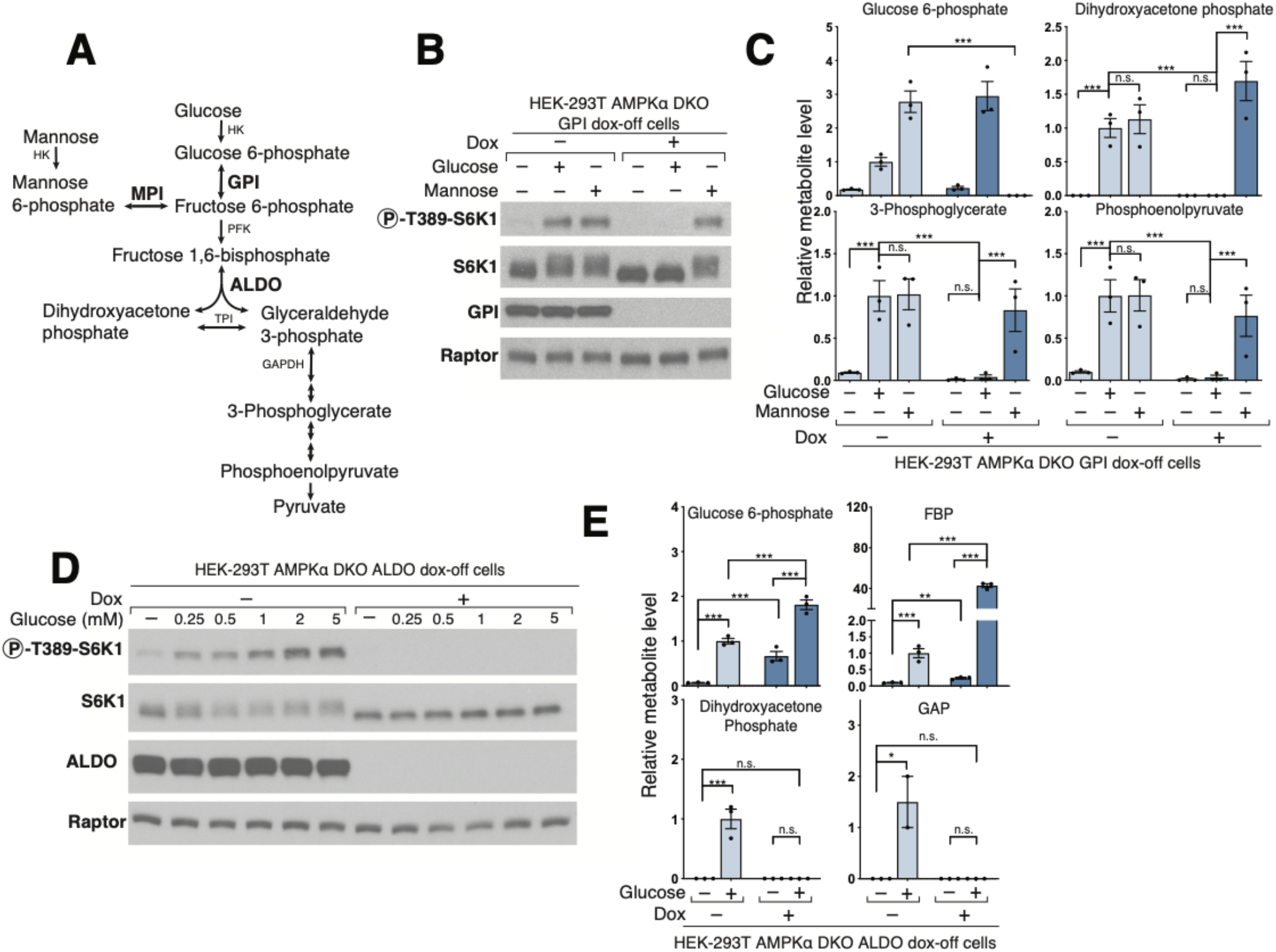
Glucose phosphoisomerase and Aldolase are upstream of the metabolite that signals glucose availability to mTORC1. **A)** Diagram of glucose and mannose metabolism emphasizing the roles of Glucose phosphoisomerase (GPI), Mannose phosphoisomerase (MPI), and Aldolase (ALDO) in glycolysis. **B)** GPI is required for glucose, but not mannose, to activate mTORC1. AMPKαDKO GPI dox-off cells were treated with doxycycline (dox) for 10 days, and placed for 2 hours in media with either no sugar (−), glucose (+Gluc), or mannose (+Mann). Cell lysates were analyzed by immunoblotting for the phosphorylation state or levels of the indicated proteins. **C)** GPI is required for the metabolism of glucose beyond glucose 6-phosphate, but not for that of mannose. Cells were treated as in (B) and metabolite extracts analyzed by LC/MS. Metabolite levels are reported relative to the −dox, +glucose condition. **D)** ALDO is required for the activation of mTORC1 by glucose. HEK-293T AMPKαDKO ALDO dox-off cells were treated with doxycycline for 5 days. Cells were then incubated in media containing the indicated concentrations of glucose for 3 hours. Cell lysates were analyzed by immunoblotting for the phosphorylation state and levels of the indicated proteins. **E)** Loss of ALDO expression leads to supraphysiological levels of Fructose 1,6-bisphosphate and prevents glucose from generating metabolites downstream of aldolase including dihydroxyacetone phosphate (DHAP) and glyceraldehyde 3-phosphate (GAP). HEK-293T AMPKαDKO ALDO dox-off cells were incubated for 3 hours without glucose (-Gluc) or with 2 mM glucose (+Gluc). Metabolite extracts were analyzed by LC/MS and metabolite levels are reported relative to the −dox, +glucose condition. Bar graphs represent mean ± SEM, n = 3 per condition, FBP, fructose 1,6-bisphosphate; GAP, glyceraldehyde 3-phosphate (* p<0.05, ** p<0.01, *** p<0.001, n.s. not significant).

Phosphofructokinase (PFK) converts F6P made by GPI into fructose 1,6-bisphosphate (FBP), then aldolase (ALDO), of which there are three paralogues (ALDOA, ALDOB, and ALDOC), cleaves it into two triose phosphates, dihydroxyacetone phosphate (DHAP) and glyceraldehyde 3-phosphate (GAP) (Fig 2A)^21^. ALDO dox-off cells were generating by targeting all three ALDOA/B/C paralogues using CRISPR/Cas9 and rescuing with a doxycycline-repressible cDNA encoding ALDOA. In ALDO dox-off cells treated with doxycycline, glucose failed to activate mTORC1 (Fig. 2D) and did not generate GAP or DHAP while accumulating levels of F6P and FBP above the level observed in normal cells (Fig 2E). These results suggest that a metabolite downstream of aldolase activates mTORC1 and eliminate F6P and FBP as candidate molecules.

The next enzyme in glycolysis, glyceraldehyde 3-phosphate dehydrogenase (GAPDH), couples the formation of NADH from NAD^+^ to the oxidation of GAP into 1,3-Bisphosphoglycerate (1,3-BPG)^21^. To probe the role of GAPDH in mTORC1 activation, we placed cells in glucose-free media and at the same time added Koningic acid (KA), a specific GAPDH inhibitor^22,23^. Interestingly, in the presence of KA, mTORC1 remained active even after cells were deprived of glucose for three hours. Metabolite profiling showed that KA creates a dam so that even upon glucose-deprivation cells maintain high levels of metabolites upstream of GAPDH, such as GAP and DHAP, while depleting downstream metabolites (Fig 3C, Fig S2A). KA treatment of glucose-starved cells lowered ATP without reversing the effects of glucose starvation on NAD+ or NADH (Fig S2A). Importantly, expression of a KA-resistant GAPDH from *T. kongii* (TK-GAPDH)^22,23^ reversed the increase in GAP and DHAP caused by KA and restored the mTORC1 inhibition normally caused by glucose starvation (Fig 3D-E, S2B). KA prevented mTORC1 inhibition only if it was added at the beginning of the glucose withdrawal period, consistent with its impacts on GAP and DHAP levels (Fig 3B-C). Doxycycline treatment of GAPDH dox-off cells had similar effects on metabolite levels and mTORC1 activity as KA, indicating that the GAPDH protein itself is not required for its inhibition to protect mTORC1 from glucose starvation (Fig 3F, S2C).

**Figure 3.**
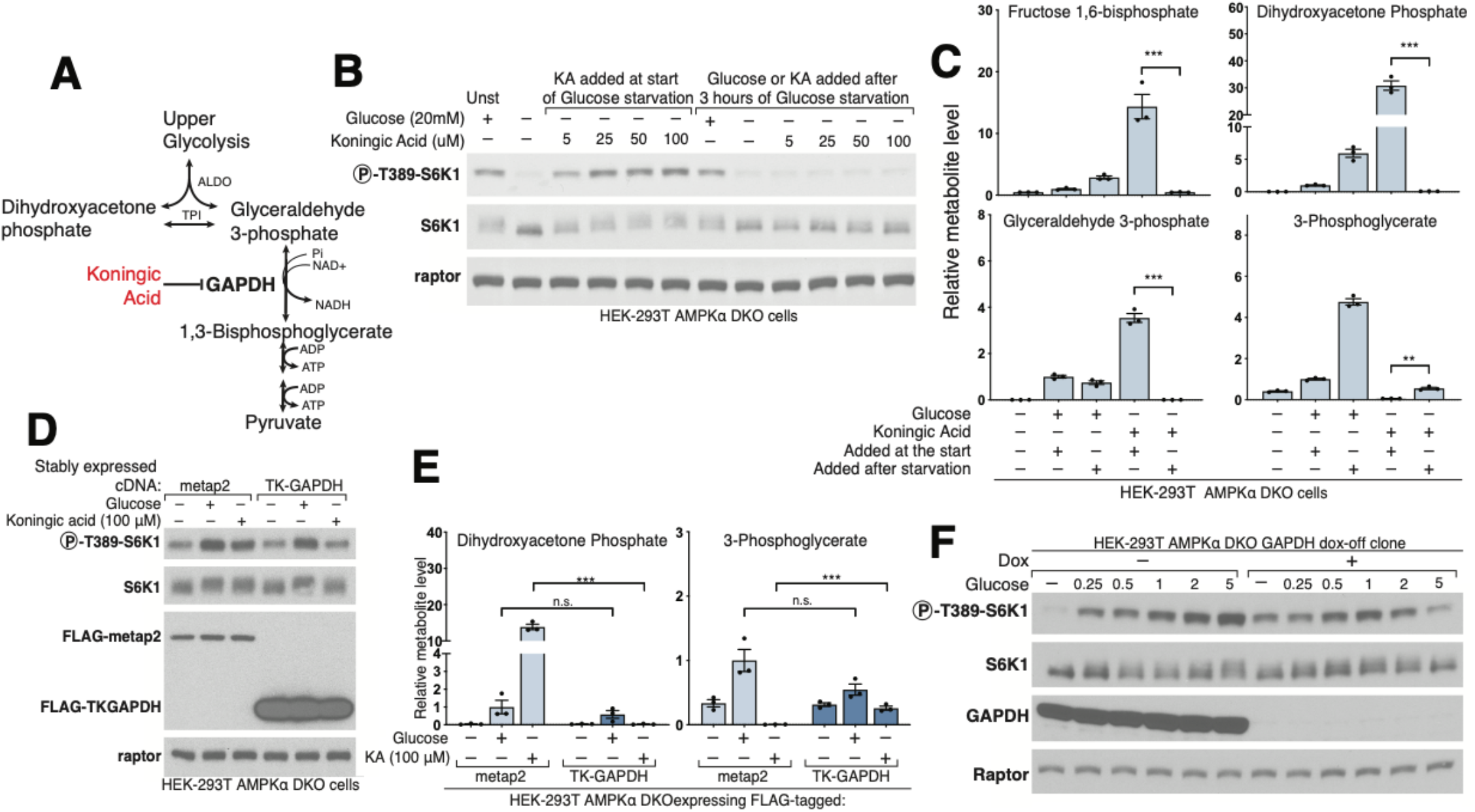
Glyceraldehyde 3-phosphate dehydrogenase (GAPDH) is downstream of the metabolite that signals glucose availability to mTORC1. **A)** Diagram of glycolysis with an emphasis on the role of GAPDH and including its inhibitor Koningic Acid. **B)** Inhibition of GAPDH by Koningic acid (KA) prevents the suppression of mTORC1 normally caused by glucose starvation, but only if KA is added at the beginning of the starvation period. AMPKαDKO HEK-293T cells were incubated with glucose or starved of it for 3 hours. Koningic acid was added to cells either at the beginning of the 3-hour glucose starvation period or for 15 minutes after a 3-hour starvation. Cell lysates were analyzed by immunoblotting for the phosphorylation state or levels of the indicated proteins. **C)** GAPDH inhibition by 50 μM KA prevents depletion of metabolites upstream of GAPDH upon glucose starvation, but only if added at the beginning of the starvation period. Cells were treated as in (B), metabolite extracts were analyzed by LC/MS, and metabolite levels are reported relative to the cells not starved of glucose. **D)** Overexpression of the KA-resistant version of GAPDH from the fungus *T. kongii* (TK-GAPDH) eliminates the effects of KA on mTORC1 signaling. Cells stably expressing FLAG-metap2 or FLAG-TK-GAPDH were incubated under the indicated conditions. Cell lysates were analyzed by immunoblotting for the phosphorylation state or levels of the indicated proteins. **E)** Overexpression of TK-GAPDH prevents the accumulation of metabolites upstream of GAPDH normally caused by KA treatment in glucose-starved cells. Cells were treated as in (D). Metabolite extracts were analyzed by LC/MS and relative metabolite levels are reported. **F)** Loss of GAPDH expression has the same phenotype as inhibition of GAPDH by KA. GAPDH dox-off cells treated with doxycycline maintain mTORC1 activity even after a 3-hour starvation of glucose. Bar graphs represent mean ± SEM, n = 3 per condition (** p<0.01, *** p<0.001, n.s. not significant).

Collectively, these data indicate that a glycolytic intermediate downstream of ALDO but upstream of GAPDH is required for glucose to activate mTORC1. Candidate molecules include GAP, DHAP, or derivatives of either. While GAP and DHAP can spontaneously interconvert, the rate of DHAP conversion to GAP is negligible in the absence of triose phosphate isomerase (TPI)^24^ (Fig. 4A), so to differentiate between the two, we generated a TPI dox-off cell line and measured DHAP and GAP levels. GAP was measured as its aniline adduct for technical reasons as explained in Figure S3. In glucose-starved cells, loss of TPI greatly slowed the catabolism of DHAP without seemingly impacting that of GAP, consistent with the direction of normal net flux being from DHAP to GAP (Fig. 4B). Given this, we reasoned that a kinetic analysis afforded a simple way to discern which molecule is most relevant to mTORC1 activation: if glucose starvation inhibited mTORC1 more slowly in cells lacking TPI, it would favor DHAP, while, if the timing of mTORC1 inhibition was unaffected, it would favor a model in which GAP has a key role. Quite revealingly, TPI loss markedly slowed the inhibition of mTORC1 caused by glucose starvation (Fig. 4C,D). Thus, TPI is downstream of the metabolite that activates mTORC1, which favors DHAP or a molecule derived from it (except GAP), as the preferred candidate.

**Figure 4.**
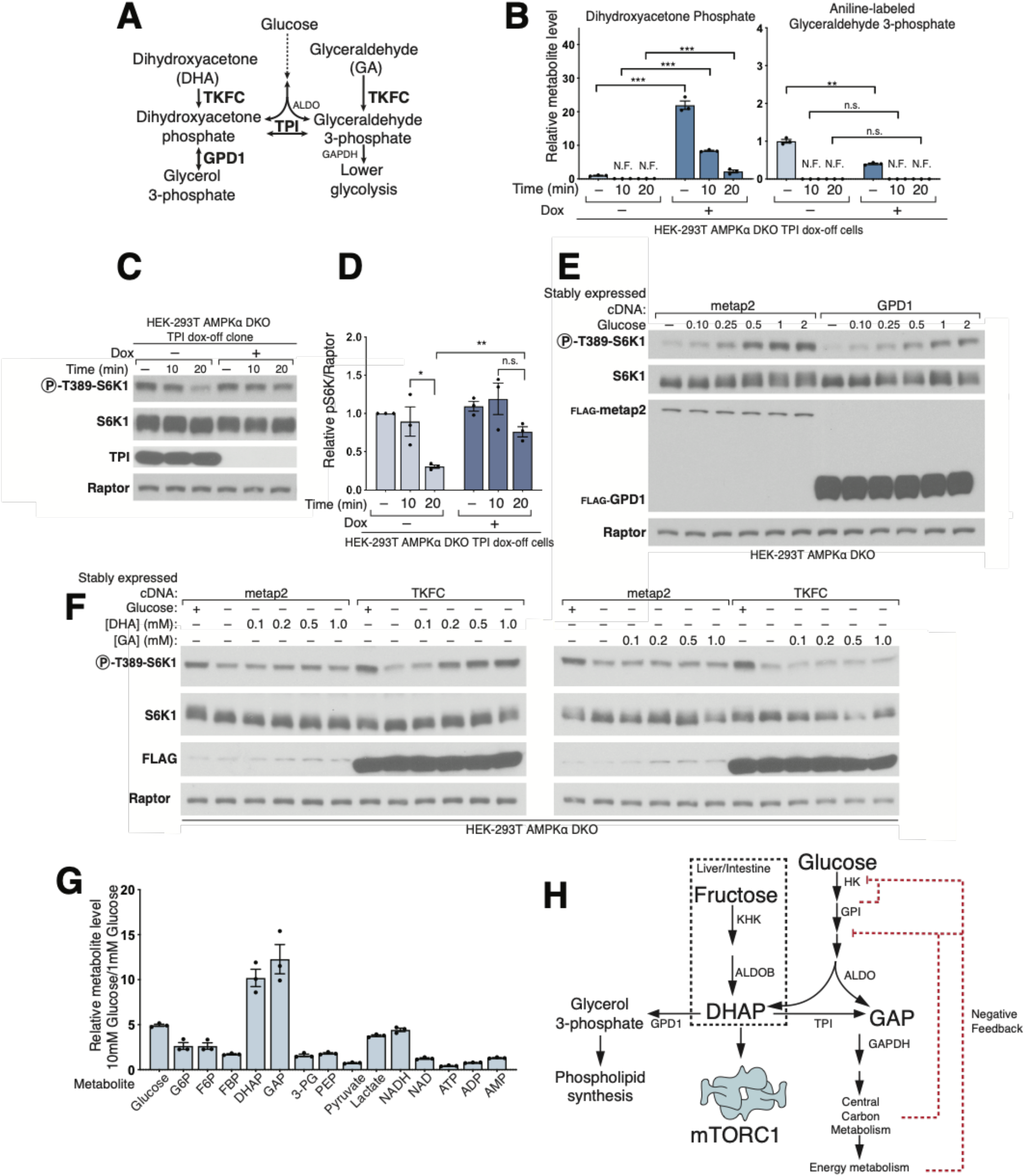
Triose phosphate isomerase (TPI) is downstream of the metabolite that signals glucose availability to mTORC1 and DHAP synthesis is sufficient to activate mTORC1. **A)** Model depicting the metabolism of trioses and triose-phosphates and emphasizing the roles of triokinase (TKFC), glycerol kinase (GK), triose phosphate isomerase (TPI), and glycerol 3-phosphate dehydrogenase (GPD1). **B)** Loss of TPI leads to supraphysiological levels of dihydroxyacetone phosphate (DHAP) and its slower catabolism following glucose starvation. The levels of glyceraldehyde 3-phosphate (GAP) are decreased but it is consumed at a normal rate. AMPKαDKO TPI dox-off HEK-293T cells were treated with doxycycline for 10 days. Cells were then starved of glucose for the indicated periods of time, and metabolite extracts were prepared and analyzed by LC/MS. **C)** TPI loss slows the inhibition of mTORC1 caused by glucose starvation. Cells were treated as in (B), and cell lysates were analyzed by immunoblotting for the phosphorylation state and levels of the indicated proteins. **D)** Quantification of three experiments (panel (C) shows results of one) reveals the slowed kinetics of mTORC1 inhibition in cells lacking TPI expression. **E)** Overexpression of GPD1 decreases the activation of mTORC1 by glucose. AMPKαDKO HEK-293T cells stably expressing the control protein FLAG-metap2 or FLAG-GPD1 were starved of glucose for 3 hours or starved of glucose and re-stimulated with the indicated concentration of glucose for 10 minutes. **F)** DHAP synthesis is sufficient to activate mTORC1 in the absence of glucose. AMPKαDKO HEK-293T cells stably expressing the control protein FLAG-metap2 or FLAG-TKFC were starved of glucose for 3 hours or starved and stimulated for 10 minutes with glucose or the indicated concentrations of dihydroxyacetone (DHA) or glyceraldehyde (GA). **G)** The fold-change in DHAP and GAP levels between cells in low (1 mM) and high (10 mM) glucose is greater than that for any other glycolytic metabolite. AMPKαDKO HEK-293T cells were starved of glucose for 3 hours and then were restimulated for 15 minutes with either 1 mM glucose or 10 mM glucose. Metabolite extracts were analyzed by LC/MS. **H)** Diagram depicting the metabolism of glucose in most cells and of fructose in the liver and intestine, emphasizing the position of DHAP in these pathways. Bar graphs represent mean ± SEM, n = 3 per condition (* p<0.05, ** p<0.01, *** p<0.001, n.s. not significant, N.F. no peak found).

One such derivative is glycerol 3-phosphate (G3P), a precursor in the synthesis of lipids, which glycerol 3-phosphate dehydrogenase (GPD1/GPD1L) generates by the reduction of DHAP (Fig 4A, S4A)^21^. Overexpression of GPD1 decreased DHAP and boosted G3P levels (Fig S5B), and, importantly, suppressed the activation of mTORC1 by glucose (Fig 4E). These results point to DHAP, but not G3P, as the glucose-derived metabolite necessary for transmitting the presence of glucose to mTORC1.

To test the sufficiency of DHAP in the activation of mTORC1 by glucose, we needed a way to restore DHAP levels in glucose-starved cells. This is challenging because negatively charged sugar-phosphates do not readily cross membranes and there is no known plasma membrane transporter for DHAP in eukaryotes. As an initial approach to overcome this issue, we generated cells expressing human glycerol kinase (GK). GK phosphorylates glycerol, which is membrane permeable, to make glycerol-3-phosphate, which glycerol 3-phosphate dehydrogenase can then oxidize into DHAP (Figure 4A, S4A)^21^. In cells starved of glucose and expressing GK, but not a control protein, glycerol partially restored mTORC1 activity (Fig S4B), which correlated with a partial restoration of DHAP levels as well as supraphysiological levels of G3P (Fig S4C).

While supporting a key role for DHAP in glucose sensing by mTORC1, we were not satisfied with the partial rescue of DHAP levels and mTORC1 activity we could obtain in the GK-expressing cells. Thus, we took advantage of human triokinase (TKFC), which can phosphorylate the membrane-permeable trioses dihydroxyacetone (DHA) and glyceraldehyde (GA) to make DHAP and GAP, respectively (Fig 4A)^21^. Expression of TKFC on its own had no effect on the regulation of mTORC1 signaling by glucose. However, in cells starved of glucose for 3 hours and expressing TKFC, but not a control protein, the addition of DHA for just ten-minutes was sufficient to reactivate mTORC1 (Fig 4F). Metabolite profiling confirmed that DHA addition leads to DHAP synthesis only in the TKFC-expressing cells and only partially restored GAP levels, consistent with our previous observation suggesting the key role of DHAP but not GAP in activation of mTORC1 (FigS5A). GA, when added at the same concentration as DHA in TKFC expressing cells, did not activate mTORC1, while at much higher doses (5-10 mM) it did (Fig. S5C). However, this effect correlated with the production of DHAP, likely from the fact that commercially available GA is contaminated with DHA (Fig. S5D). Lastly, DHA also restored mTORC1 activity in TPI and ALDO deficient cells expressing TKFC (Fig. S5E). Therefore, we conclude that DHAP synthesis is sufficient to activate mTORC1.

Many upstream inputs regulate mTORC1 activity, including the nutrient-sensing pathway anchored by the heterodimeric Rag GTPases that recruit mTORC1 to the lysosomal surface, its site of activation. The Rag GTPases and their positive (GATOR2, Ragulator, SLC38A9, FLCN-FNIP) and negative (GATOR1, KICSTOR) upstream regulators play key roles in the sensing of amino acids by mTORC1^1^. In addition, several stress-sensing pathways activate the ATF4 transcription factor, which suppresses mTORC1 signaling, at least in part, by upregulating the Sestrin2 leucine sensor that acts through GATOR2^25^. We found that neither loss of ATF4 nor its upstream activator, GCN2, impacted glucose sensing in AMPK DKO cells (Fig S6A-B). Recently, the Wnt pathway component Axin1 has been implicated in glucose sensing upstream of AMPK via an interaction with the Ragulator complex^15^. In our cells, loss of Axin1 did not affect the capacity of mTORC1 or AMPK to be regulated by glucose and we could not detect Ragulator in Axin1-immunoprecipitates, suggesting that the role of Axin in detecting glucose might be cell-type dependent (Fig S6C-D). In contrast to these negative results and consistent with previous work implicating the Rag GTPases in the AMPK-independent sensing of glucose by mTORC1^9,10^, loss of core components of GATOR1 or KICSTOR eliminated the capacity of glucose starvation to inhibit mTORC1 (Fig S7A-C). While loss of FLCN did not affect the ability of glucose to regulate mTORC1 (Fig S7D), that of a GATOR2 component (Mios) prevents full activation of mTORC1 by glucose (Fig S7E). Moreover, in cells overexpressing all five components of GATOR2 (WDR59, WDR24, MIOS, SEH1L, and SEC13) glucose starvation no longer inhibited mTORC1 (Fig S7F). These data suggest that DHAP signals glucose availability to mTORC1 via the Rag GTPase pathway and specifically the GATOR2-GATOR1-KICSTOR input. Consistent with previous reports^9^ and the known roles of the Rag GTPases in mTORC1 activation, starvation of glucose regulated the localization of mTORC1 to the lysosomal surface, albeit to a lesser degree than that of amino acids (Fig S7G-H).

Our extensive attempts to identify DHAP sensor proteins akin to Sestrin2 for leucine^26^ and CASTOR1 for arginine^27^ have been unsuccessful. In addition, our existing data suggest that none of the enzymes known to consume or generate DHAP are also a DHAP sensor for the mTORC1 pathway. TPI cannot be the sensor because while its loss slows inhibition of mTORC1 upon glucose starvation, ultimately it is not necessary for glucose to regulate the pathway (Fig 4B, S5E). Likewise, none of the aldolase paralogues can be the sensor because DHA treatment of TKFC-expressing cells activates mTORC1 in cells lacking all of them (Fig S5E). Lastly, loss of GPD1 and its paralogue GPD1L does not prevent glucose regulation of mTORC1, ruling them out as sensors (Fig S5F). Therefore, we conclude that the DHAP sensor is likely not a protein currently known to interact with DHAP. GAPDH was previously reported to bind the mTORC1-activating Rheb GTPase and to be a GAP sensor for the mTORC1 pathway^16^, but in our hands we have not detected an interaction between GAPDH and Rheb (Fig S6E). Similarly, ALDOA was recently proposed to act as a glucose sensor to regulate the AMPK pathway via a nutrient regulated interaction with the v-ATPase complex^15^; however, we are unable to detect an interaction between ALDOA and the v-ATPase component ATP6V1B2 in our cell system (Fig S6E).

## Discussion

While confirmation that DHAP is a *bona fide* signaling molecule awaits the discovery and manipulation of the sensing mechanism, our current findings support the conclusion that it has a key role in transmitting glucose availability to mTORC1: (i) the synthesis of DHAP is sufficient to activate mTORC1 in the absence of glucose; (ii) enzymes upstream of DHAP are necessary for the activation of mTORC1 by glucose (iii) and those downstream for its suppression by glucose starvation.

An important question is why the mTORC1 pathway acquired the capacity to respond to glucose in a manner independent of the cellular energy status. Since mTORC1 increases the rate of glycolysis by regulating HIF1α^28^, it makes sense for it to sense glucose availability. However this doesn’t answer why, in particular, the cell uses DHAP to do so. In retrospect, there are many reasons why DHAP is particularly well suited to play a role as a signaling molecule. We compared the glycolytic metabolite levels of cells in media containing 10 or 1 mM glucose (Fig 4G). DHAP, along with GAP, change about 10-fold between these two glucose concentrations, the most of any other metabolite in glycolysis. The dynamic nature of DHAP makes it well suited to signal glucose availability. Of note, cellular ATP or AMP levels are not nearly as dynamic (Fig 4G), which reflects the involvement of other nutrients besides glucose, such as glutamine and fatty acids, in energy metabolism (Fig 4G). Second, DHAP, in addition to being a glycolytic intermediate, is a precursor for the glycerol backbone used in the synthesis of triglycerides and phospholipid synthesis^21^, processes that mTORC1 promotes when active^29^. Linking DHAP levels to mTORC1 activation ensures that lipid synthesis is only fully activated when the precursor metabolite DHAP is at acceptable levels. Therefore, DHAP sensing by the mTORC1 pathway could play a critical role in the post-prandial *de novo* lipogenesis in adipose tissue. In agreement with this notion, a previous study demonstrated that GLUT4 overexpression in adipose tissue led to greater increases in insulin stimulated mTORC1 activation suggesting that glucose uptake, in concert with insulin action, plays a key role in driving mTORC1-dependent anabolism^30^. Third, because glucose and fructose differ in how they lead to DHAP synthesis, DHAP sensing may allow the mTORC1 pathway to respond with different potency to these dietary sugars, a feat that is harder to accomplish for AMPK. Specifically, fructose requires only two steps to make DHAP but glucose requires four and fructose metabolism to DHAP bypasses the steps of glycolysis that are subject to feedback inhibition^31^, suggesting that the rate of fructose conversion to DHAP is likely to be less restrained relative to that from glucose and therefore could be a more potent activator of mTORC1 (Fig 4H). And, importantly, while many different cell types metabolize glucose, fructose is primarily metabolized by the small intestine and the liver^32^. In the latter, fructose stimulates lipogenesis and drives diet-induced fatty liver disease^33^. While much more work is necessary to determine the role of DHAP sensing by mTORC1 in vivo, it is tempting to speculate that the link between dietary fructose and lipogenesis might involve the activation of anabolic processes by mTORC1. Future work should also clarify the role in mTORC1 activation of other carbon sources such as lactate or glycerol, which should have the potential to activate mTORC1 by way of DHAP production under the right physiological contexts.

**Supplemental Figure 1.**
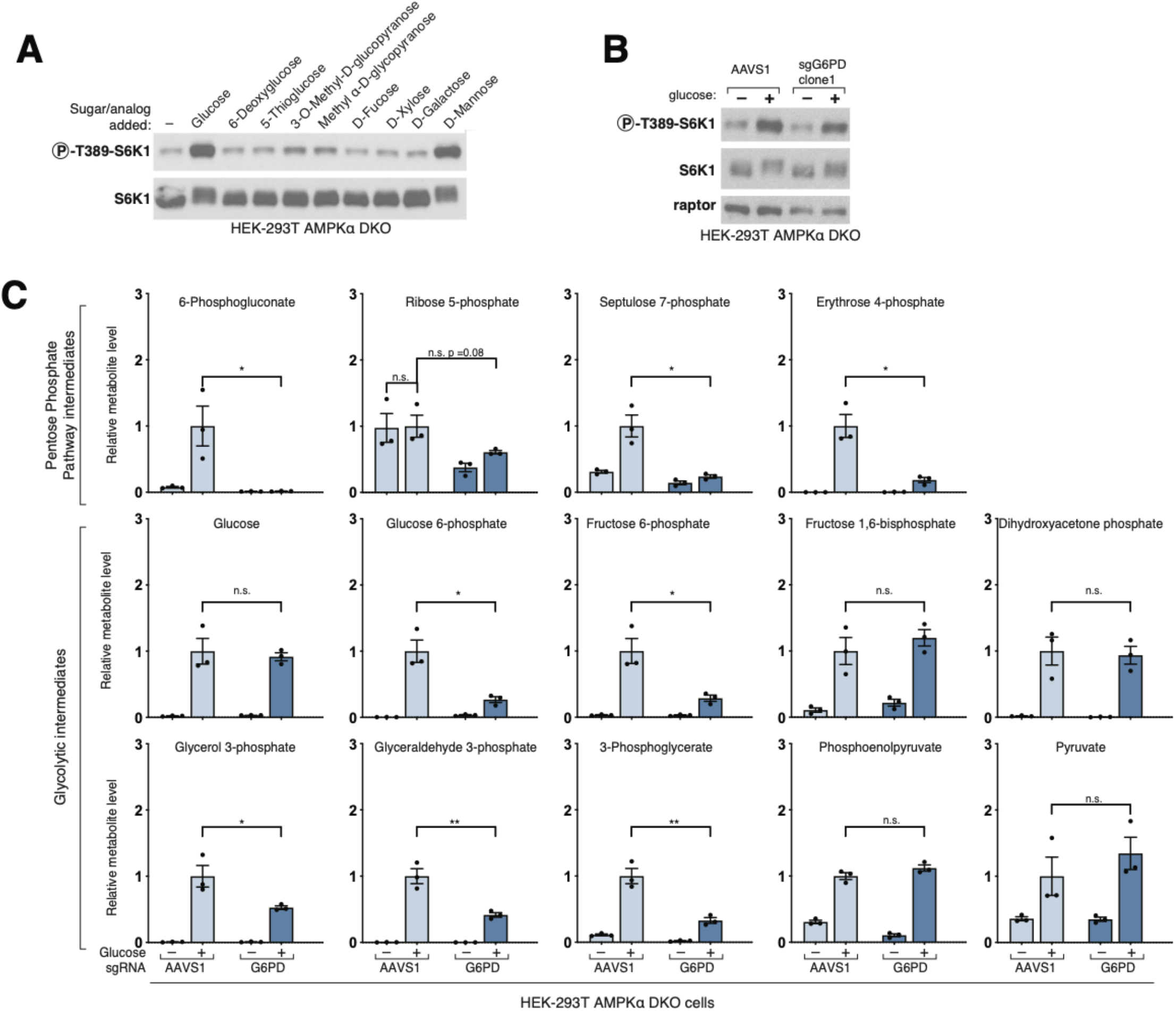
Glucose metabolism via the pentose phosphate pathway (PPP) is not required for the activation of mTORC1. **A)** AMPK DKO cells were incubated for 1 hour without glucose, then either maintained without an added sugar or the indicated sugar or sugar analog was added for 15 minutes prior to cell lysis. Whole cell lysates were analyzed for the phosphorylation state or level of S6K1 by immunoblotting. **B)** Cells targeted with CRISPR/Cas9 using a control guide (AAVS1) or one against G6PD were starved of glucose for 1 hour or starved for 50 minutes and re-stimulated with glucose for 10 minutes. Whole cell lysates were analyzed as in (A). Loss of G6PD did not prevent the mTORC1 pathway from responding to glucose starvation and restimulation. **C)** Loss of G6PD caused a predictable decrease in pentose phosphate intermediates but had no impact on most glycolytic intermediates. Cells were treated the same as in (B and metabolite extracts were analyzed by LC/MS for the indicated metabolites. Bar graphs represent mean ± SEM, n = 3 per condition (* p<0.05, ** p<0.01, n.s. not significant).

**Supplemental Figure 2.**
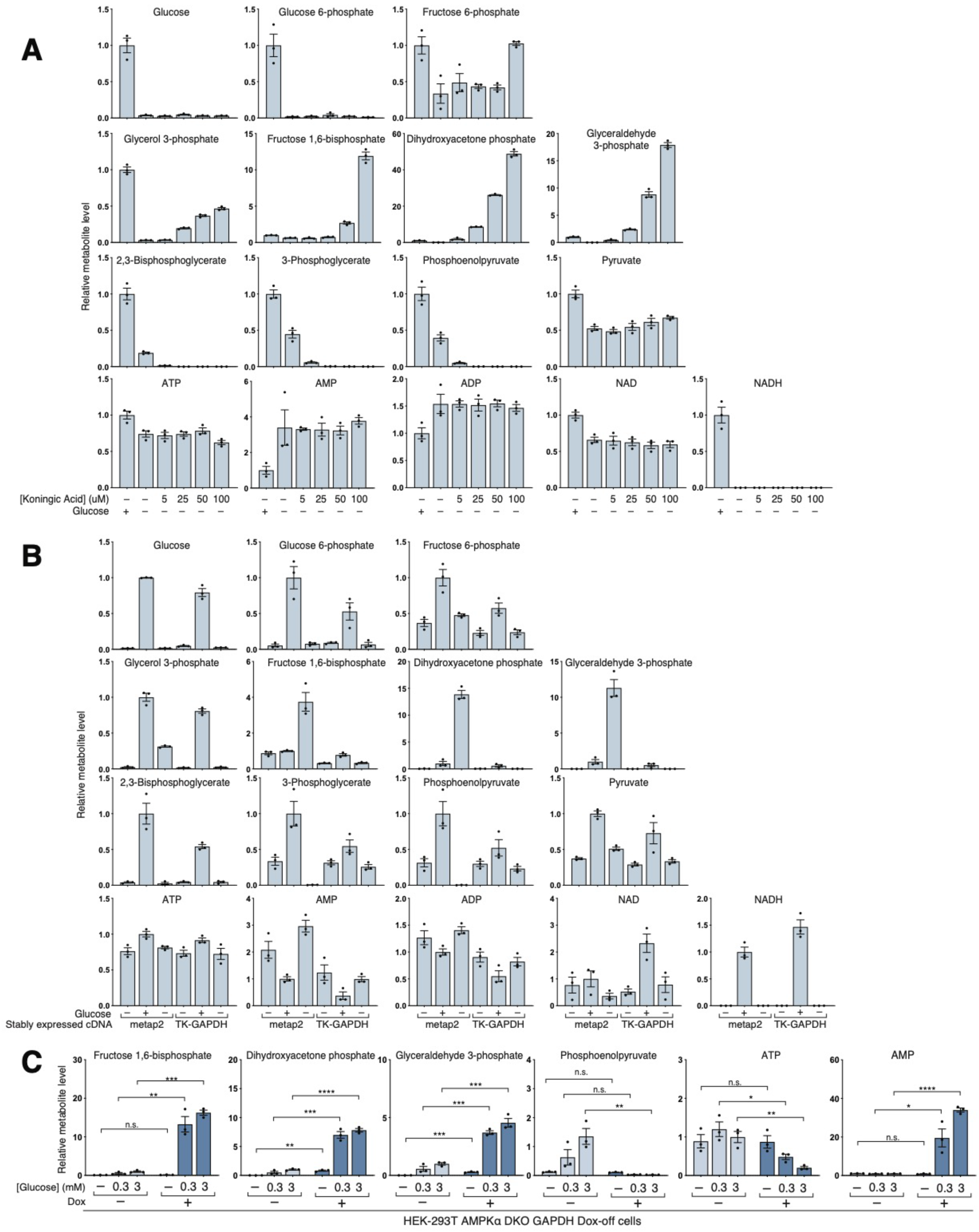
Metabolite profiling of cells lacking GAPDH activity. **A)** In a dose-dependent manner, KA leads to the increase of the metabolites upstream of GAPDH despite the lack of glucose in the media. Metabolites downstream of GAPDH are predictably depleted upon KA treatment. NAD levels fall slightly upon glucose starvation and are not affected by KA treatment. NADH became undetectable in the absence of glucose, consistent with the role of glucose metabolism in NADH production. The effects of glucose deprivation on ATP/ADP/AMP levels was not reversed by KA treatment. HEK-293T cells were treated with the indicated concentrations of Koningic Acid (KA) at the beginning of a glucose starvation period, and metabolites were extracted and analyzed by LC/MS. **B)** Overexpression of TK-GAPDH prevents all the effects on glycolytic metabolites caused by KA. Cells stably expressing FLAG-metap2 or FLAG-TK-GAPDH were incubated in the indicated conditions, metabolites were extracted and analyzed by LC/MS. **C)** Consistent with pharmacological inhibition of GAPDH, loss of GAPDH expression increases levels of GAP and DHAP and decreases levels of PEP. GAPDH Dox-off cells were exposed to the indicated glucose concentrations in the absence or presence of dox. Bar graphs represent mean ± SEM, n = 3 per condition (* p<0.05, ** p<0.01, *** p<0.001, **** p<0.0001, n.s. not significant).

**Supplemental Figure 3.**
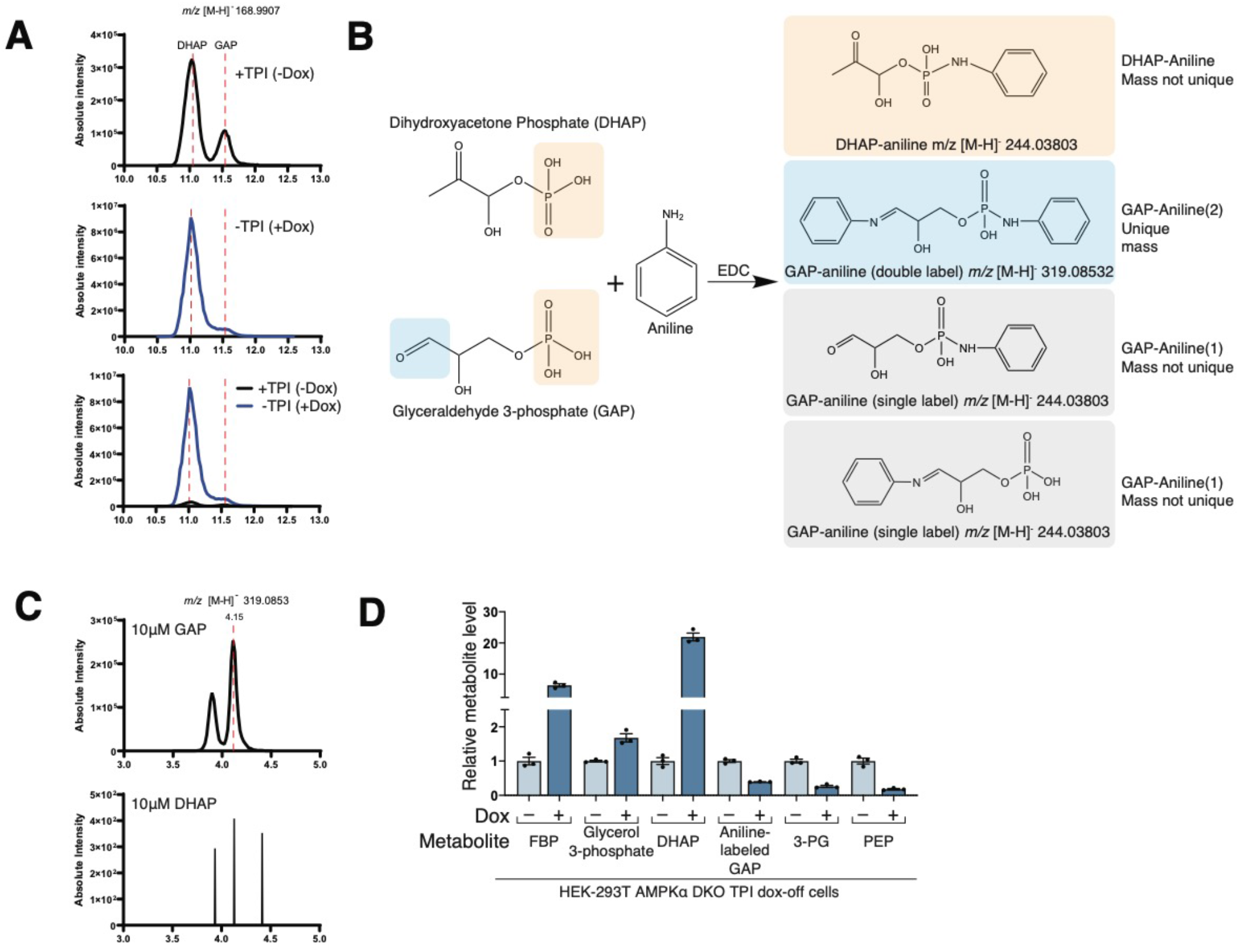
GAP can be measured as its aniline adduct as it has a unique mass. **A)** Normally the DHAP and GAP metabolites elute at similar retention times (11 mins vs 11.5 mins, respectively) but the peaks are separate enough to integrate independently. However, when cells lose TPI expression, the DHAP levels increase by an order of magnitude, and the peak normally integrated to obtain the quantities of GAP is obscured by the broader DHAP peak with a shoulder that extends into the normally eluting GAP peak at 11.5 mins. Because the mass of DHAP and GAP are the same, quantifying them accurately in the context of TPI loss is an analytical challenge. HEK-293T AMPK DKO TPI dox-off cells were treated with (+Dox) or without dox (-Dox) for 10 days. Then the media was replaced with fresh RPMI media containing 5 mM glucose for thirty minutes. Metabolites were extracted and analyzed by HILIC LC/MS. Peak traces are shown for the +TPI (-Dox) condition in black and −TPI (+Dox) in blue both separately, with different y-axis scales, or overlaid to highlight the observed changes in the DHAP. **B)** DHAP and GAP can form aniline adducts that are singly labeled with an *m/z* value of 244.03803 but only GAP can form an aniline adduct that is doubly labeled with an *m/z* value of 319.08532. **C)** As expected, only GAP formed a product with *m/z* value of 319.0853 when reacted with aniline. 10 μM GAP or DHAP standards were reacted with aniline in the presence of EDC in 80% methanol to yield their respective aniline adducts. The reaction was analyzed by C8 LC/MS. **D)** Metabolites upstream of TPI increase and metabolites downstream of TPI decrease in the absence of TPI expression. If GAP levels are measured from the shoulder of the DHAP peak in the normal HILIC LC/MS method (red bars) it paradoxically appears to also increase upon loss of TPI expression. However, if we integrate the GAP-aniline(2) peak, it then follows the expected pattern of decreasing upon loss of TPI, similar to the pattern observed for 3-phophsoglycerate (3-PG) and phosphoenolpyruvate (PEP). HEK-293T AMPK DKO TPI dox-off cells were treated as in (A) and metabolites were extracted in 80% methanol. Half of the extract was used in an aniline labeling reaction and analyzed by C8 LC/MS and the other half was analyzed by HILIC LC/MS. Bar graphs represent mean ± SEM, n = 3 per condition.

**Supplemental Figure 4.**
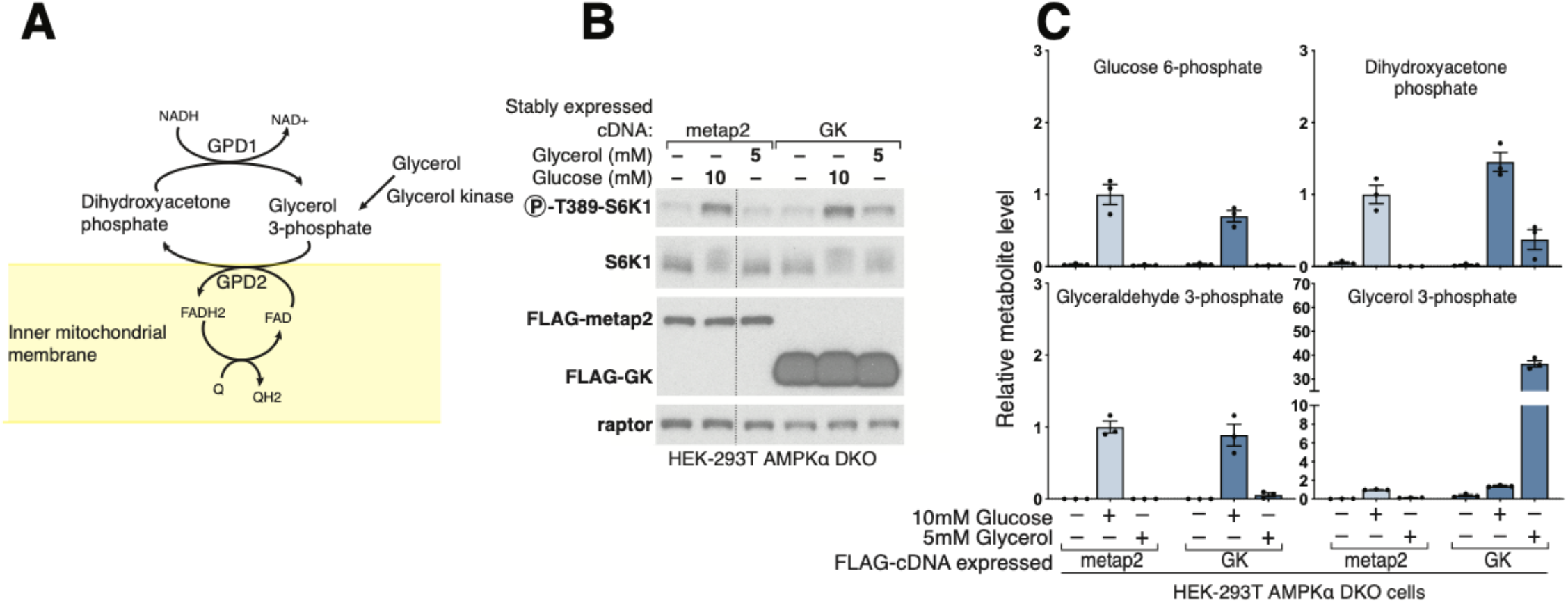
Glycerol Kinase-mediated glyceraldehyde 3-phosphate (G3P) synthesis partially activates mTORC1. **A)** Model detailing the metabolism of glycerol, and its connection to the G3P-shuttle and the glycolytic metabolite dihydroxyacetone phosphate (DHAP). The role of GPD1 in coupling DHAP reduction to NADH oxidation is emphasized. **B)** HEK-293T AMPK DKO cells stably overexpressing either a FLAG-tagged control cDNA (metap2) or glycerol kinase (GK) were incubated with the indicated glucose or glycerol concentrations for 1 hour. Whole cell protein lysates were analyzed by immunoblotting for the phosphorylation state and levels of the indicated proteins. **C)** Cells were treated as in (B). Metabolite extracts were analyzed by LC/MS. Notably, glycerol addition to cells expressing GK, but not the control protein, generated supraphysiological levels of G3P but only partially restored DHAP levels. Dashed line on panel B represents the deletion of irrelevant intervening lanes in the western blot. Bar graphs represent mean ± SEM, n = 3 per condition.

**Supplemental Figure 5.**
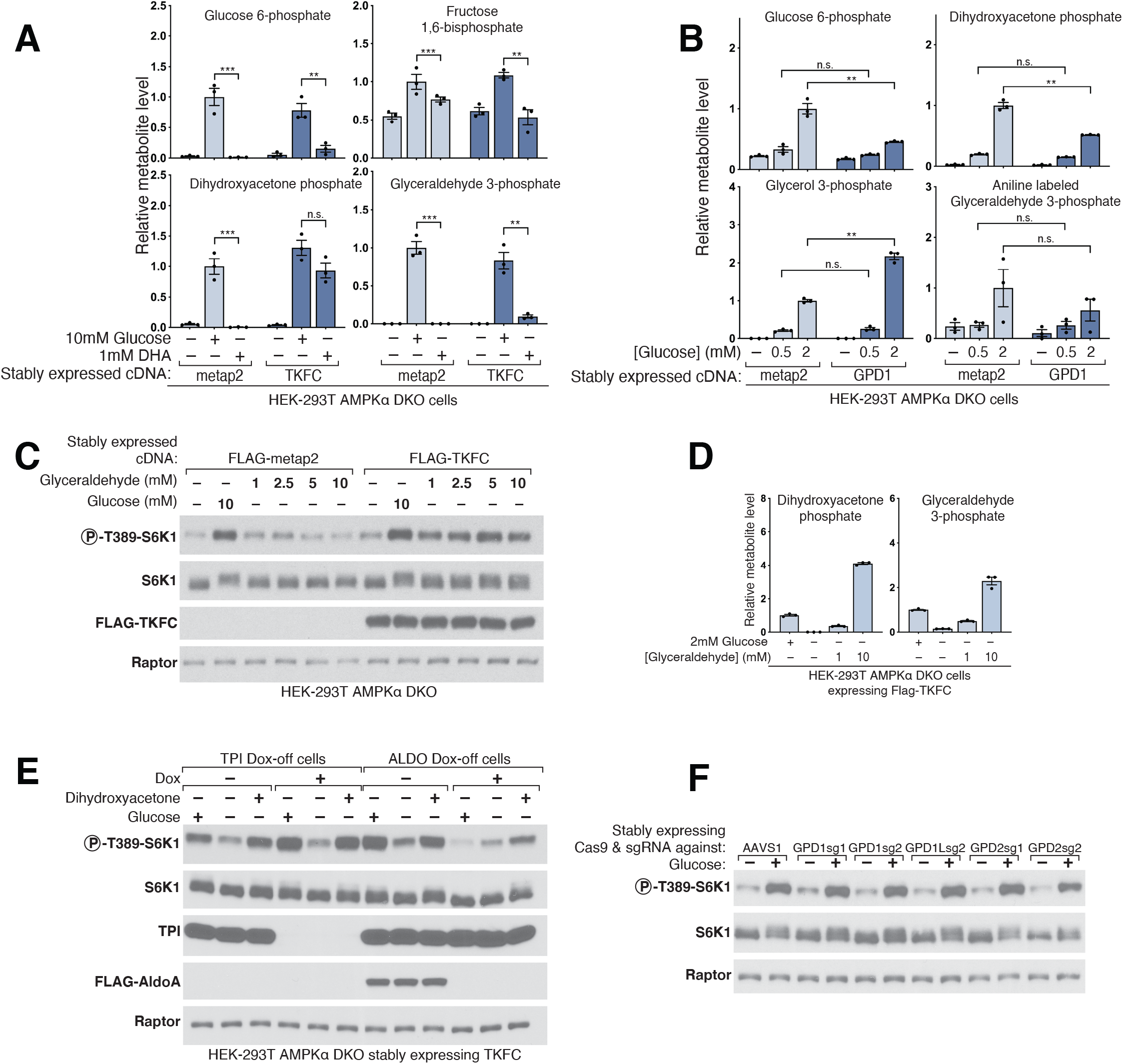
DHAP plays a key role in the activation of mTORC1 by glucose. **A)** Dihydroxyacetone (DHA) makes DHAP only in TKFC-expressing cells, restoring levels to nearly those seen upon glucose restimulation. Moreover, DHA treatment has little to no effect on glucose 6-phosphate (G6P) or fructose 1,6-bisphosphate (FBP) levels and only partially rescues glyceraldehyde 3-phosphate (GAP) levels. HEK-293T AMPK DKO cells expressing a FLAG-tagged control cDNA (metap2) or triose kinase (TKFC) were incubated with the indicated concentrations of glucose or DHA for 15 minutes following a three-hour glucose starvation. Metabolite extracts were analyzed by LC/MS. **B)** GPD1 overexpression decreases the levels of several glycolytic intermediates (G6P and DHAP are shown) but increases those of glycerol 3-phosphate. HEK-293T AMPK DKO cells expressing a FLAG-tagged control cDNA (metap2) or glycerol 3-phosphate dehydrogenase (GPD1) were incubated with the indicated glucose concentration for 15 mins following a 3-hour glucose starvation. Metabolite extracts were analyzed by HILIC LC/MS for G6P, DHAP, and glycerol 3-phosphate, or aniline labeled and analyzed by C8 LC/MS for GAP-aniline. **C)** Glyceraldehyde (GA), when given at concentrations above 5 mM, partially rescued mTORC1 signaling in the absence of glucose. HEK-293T AMPK DKO cells expressing a FLAG-tagged control cDNA (metap2) or triose kinase (TKFC) were incubated with the indicated concentrations of glucose or GA for one hour. Whole cell lysates were analyzed for the phosphorylation state and levels of the indicated proteins by immunoblot. **D)** The concentration required to obtain partial rescue with GA are capable of making DHAP along with GAP, here measured as the derivative GAP-aniline. Cells were treated as in (C). Metabolite extracts were analyzed by HILIC LC/MS for DHAP or reacted with aniline and analyzed by C8 LC/MS for GAP-aniline. **E)** DHA does not require TPI or ALDO expression in order to activate mTORC1. TPI Dox-off and ALDO Dox-off cells were treated with either glucose or DHA in the absence or presence of doxycycline, cell lysates were analyzed by immunoblotting for the phosphorylation state and levels of the indicated proteins. **F)** Loss of GPD1 or GPD1L does not affect glucose sensing by mTORC1. HEK-293T AMPK DKO stably expressing Cas9 with either a control guide (AAVS1) or guides against GPD1, GPD1L, or GPD2 were starved of glucose for 3 hours or starved for 3 hours and restimulated with glucose for 15 minutes. Whole cell lysates were analyzed for the phosphorylation and levels of the indicated proteins by immunoblot. Bar graphs represent mean ± SEM, n = 3 per condition (* p<0.05, ** p<0.01, *** p<0.001, n.s. not significant).

**Supplemental Figure 6.**
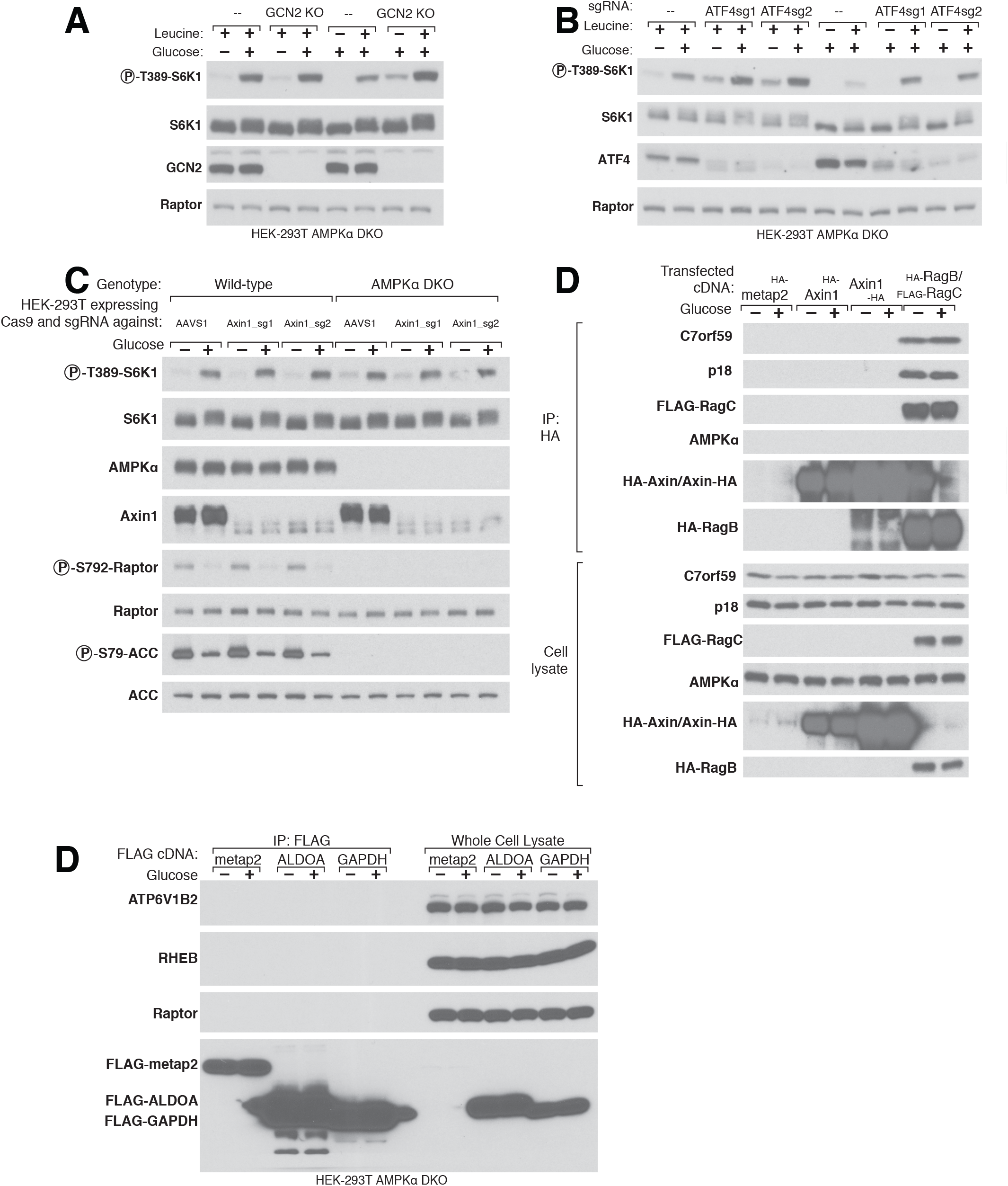
Glucose sensing does not require canonical stress pathway components. **A)** Loss of GCN2 changed baseline mTORC1 activity but did not affect its ability to sense glucose or leucine starvation. HEK-293T AMPK DKO or AMPK/GNC2 triple KOs (TKO) were starved of glucose or leucine for 3 hours or starved and restimulated with glucose or leucine for 15 minutes. Whole cell lysates were analyzed for the phosphorylation and levels of the indicated proteins by immunoblot. **B)** While loss of ATF4 increased baseline mTORC1 activity, it did not change the ability to sense glucose or leucine starvation. HEK-293T AMPK DKO stably expressing Cas9 with either a control guide (AAVS1) or guides against ATF4 were starved of glucose or leucine for 3 hours or starved and restimulated with glucose or leucine for 15 minutes. Whole cell lysates were analyzed for the phosphorylation and levels of the indicated proteins by immunoblot. **C)** Loss of Axin1 does not affect glucose sensing by the mTORC1 pathway or that of AMPK. Wildtype or AMPK DKO HEK-293T cells stably expressing Cas9 and either a guide against a control locus (AAVS1) or Axin1 were starved of glucose or starved and restimulated with glucose. Whole cell lysates were analyzed for the phosphorylation state and levels of the indicated proteins by immunoblotting. **D)** RagC/RagB but not Axin1 was able to co-immunoprecipitate the Ragulator components c7orf59 and p18. None of the tested proteins were able to co-immunoprecipitate AMPK. HEK-293T AMPK DKO transfected with cDNAs for either HA-tagged metap2, Axin1(N-terminal or C-terminal tags), or HA-RagB/Flag-RagC were starved of glucose or starved and restimulated with glucose for 15 minutes. HA-immunopreciptates and whole cell lysates were analyzed for the levels of the indicated proteins by immunoblotting. **E)** ALDOA did not co-immunopreciptate the v-ATPase component ATP6V1B2 nor did GAPDH co-immunopreciptate RHEB. HEK-293T AMPK DKO stably expressing FLAG-metap2, FLAG-ALDOA, or FLAG-GAPDH were starved of glucose or starved and restimulated with glucose. FLAG-IPs or whole cell lysates were analyzed for the levels of the indicated proteins by immunoblotting.

**Supplemental Figure 7.**
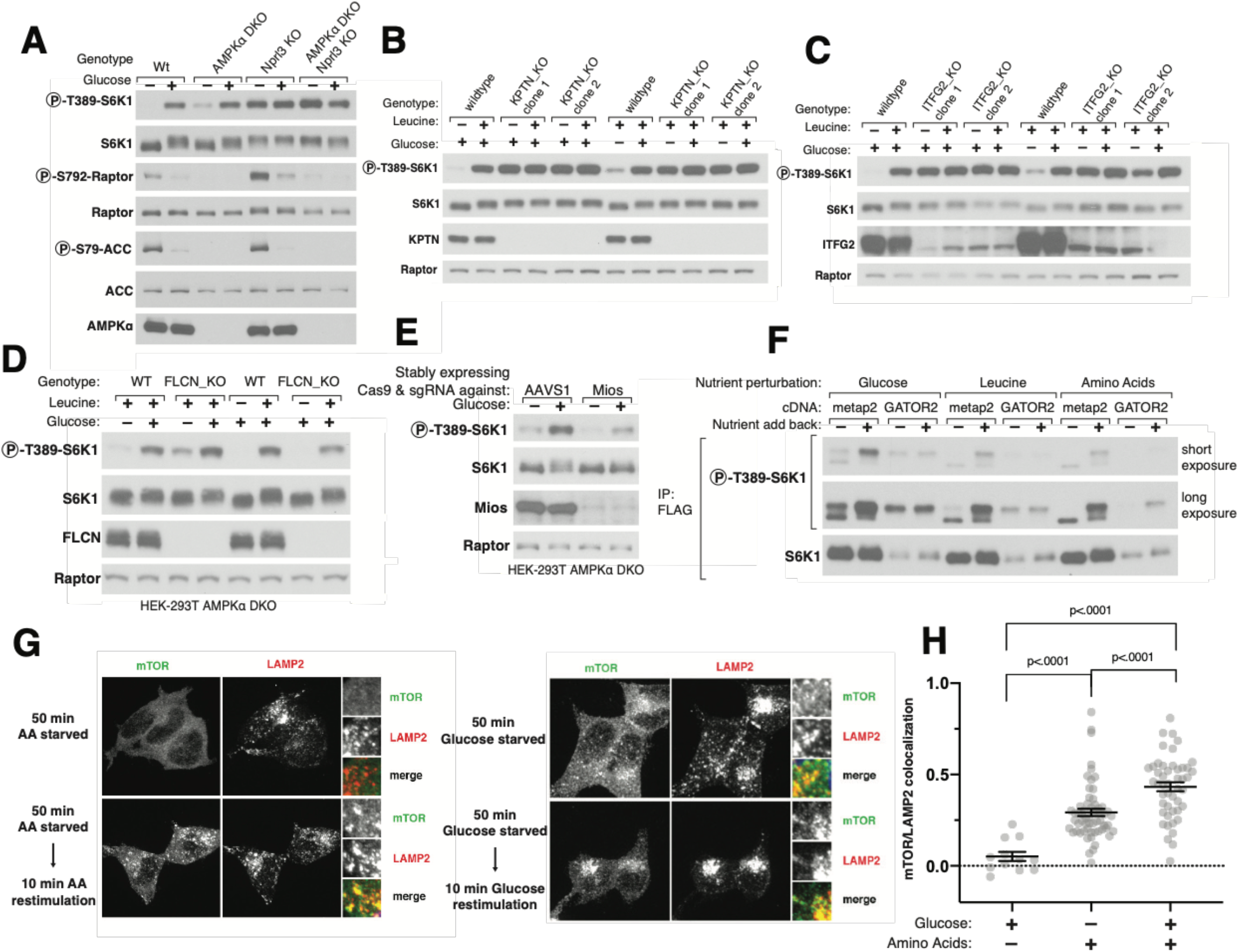
Glucose sensing by mTORC1 requires an intact GATOR/KICSTOR pathway. **A)** In Nprl3 KO cells, independent of the AMPK status, mTORC1 was insensitive to glucose starvation. HEK-293T wildtype, AMPK DKO, Nprl3 KO, or AMPK/Nprl3 TKO cells were starved of glucose or starved and restimulated with glucose for 15 minutes. Whole cell lysates were analyzed for the phosphorylation state or levels of the indicated proteins. **B-C)** Loss of either of the two components of KICSTOR prevents mTORC1 from sensing glucose or leucine. Wildtype HEK-29T cells or KOs for the KICSTOR complex genes KPTN (panel B) or ITFG2 (panel C) were starved of either glucose or leucine or starved and restimulated with glucose or leucine. Whole cell lysates were analyzed for the phosphorylation state or level of the indicated proteins. **D)** Loss of FLCN does not prevent mTORC1 from sensing glucose or leucine. HEK-293T AMPK DKO or AMPK/FLCN TKO cells were starved of either glucose or leucine or starved and restimulated with glucose or leucine. Whole cell lysates were analyzed for the phosphorylation state and level of the indicated proteins. **E)** A decrease in Mios expression leads to a concomitant decrease in the activation by glucose of mTORC1. HEK-293T AMPK DKO stably expressing Cas9 and either a control guide or one targeting the GATOR2 component Mios were starved of glucose or starved and restimulated with glucose. Whole cell lysates were analyzed for the phosphorylation state and levels of the indicated proteins by immunoblotting. **F)** Overexpression of the five components of GATOR2 prevents glucose or leucine starvation from regulating mTORC1. However, full amino acid starvation still regulated mTORC1 activity. HEK-293T AMPK DKO co-transfected with cDNAs for FLAG-S6K and either metap2 or all five components of the GATOR2 complex (Mios, WDR24, WDR59, SEC13, SEH1L) were starved of glucose, leucine, or amino acids or were starve and restimulated with glucose, leucine, or amino acids. FLAG-IP or whole cell lysates were analyzed for the phosphorylation state or levels of the indicated proteins. **G)** As previously reported, both glucose and amino acids regulate mTOR localization to the lysosome consistent with glucose regulating the Rag-GTPase pathway upstream of mTORC1. AMPK DKO cells were starved of amino acids, glucose or left unstarved. Cells were fixed and permeabilized and analyzed in an immunofluorescence assay with mTOR and LAMP2 antibodies. **H)** Glucose starvation had a weaker effect on the localization of mTOR on the lysosome than starvation of all amino acids. HEK-293T AMPK DKO cells were incubated in media lacking amino acids or glucose or full RPMI media for 1 hour. Cells were fixed and permeabilized and analyzed in an immunofluorescence assay with mTOR and LAMP2 antibodies. Line and bracket indicate mean ± SEM, n≥ 12.

## Methods

### Antibodies

| Antibody Target | Company | Catalogue number |
| --- | --- | --- |
| phospho-T389-S6K1 | Cell Signaling Technology | 9234 |
| S6K1 | Cell Signaling Technology | 2708 |
| Phospho-S79-ACC | Cell Signaling Technology | 3661 |
| ACC | Cell Signaling Technology | 3662 |
| AMPKα | Cell Signaling Technology | 2532 |
| Raptor | Millipore Sigma | 09-217 |
| HA | Cell Signaling Technology | 3724 |
| FLAG | Cell Signaling Technology | 14793 |
| GPI | ABCAM | ab167394 |
| ALDOA | Cell Signaling Technology | 8060 |
| TPI | Novus Biologicals | NBP1-31470 |
| GAPDH | Cell Signaling Technology | 2118 |
| Phospho-S792-Raptor | Cell Signaling Technology | 2083 |
| GCN2 | Cell Signaling Technology | 3302 |
| Axin1 | Cell Signaling Technology | 2087 |
| C7orf59 | Cell Signaling Technology | 12284 |
| P18 | Cell Signaling Technology | 8975 |
| KPTN | ProteinTech | 16094-1AP |
| ITFG2 | Santa Cruz | Sc-134686 |
| FLCN | Cell Signaling Technology | 3697 |
| Mios | Cell Signaling Technology | 13557 |
| ATF4 | Cell Signaling Technology | 11815 |
| ATP6V1B2 | Cell Signaling Technology | 14488 |
| RHEB | Cell Signaling Technology | 4935 |
| Anti-rabbit IgG, HRP-linked | Cell Signaling Technology | 7074 |
| mTOR | Cell Signaling Technology | 2983 |
| LAMP2 | Santa Cruz | H4B4 |

### Chemicals

| Chemical name | Company | Catalogue number |
| --- | --- | --- |
| 3-O-Methyl-D-glucopyranose | Millipore Sigma | M4879 |
| methyl α-D-glycopyranoside | Millipore Sigma | M9376 |
| 5-Thio-D-glucose | Santa Cruz | sc-221044A |
| 6-Deoxy-D-glucose | Millipore Sigma | D9761 |
| 2-Deoxy-D-glucose | Millipore Sigma | D8375 |
| D-(+)-Mannose | Millipore Sigma | M6020 |
| D-(+)-Glucosamine | Millipore Sigma | G4875 |
| Glycerol | Millipore Sigma | G2025 |
| D-(+)-Glucose | Millipore Sigma | G7021 |
| D-(-)-Fructose | Millipore Sigma | F3510 |
| D-(+)-Galactose | Millipore Sigma | G0750 |
| Koningic Acid (Heptelidic Acid) | Abcam | ab144269 |
| Aniline-HCl | Millipore Sigma | A8524 |
| N-(3-Dimethylaminopropyl)-N'-ethylcarbodiimide hydrochloride (EDC) | Millipore Sigma | E6383 |
| Triethylamine | Millipore Sigma | 471283 |

### Other materials

Anti-FLAG M2 affinity gel from Millipore Sigma; XtremeGene9 and Complete Protease Cocktail from Roche; Alexa 488, 568 and 647-conjugated secondary antibodies, and Inactivated Fetal Bovine Serum (IFS) from Invitrogen; leucine-free and amino acid-free RPMI from US Biologicals; and anti-HA magnetic beads, -glucose RPMI (cat # 11879020) from ThermoFisher Scientific. DMEM high glucose (catalog number: 11995040) and DMEM low glucose (catalogue number: 11885092) were purchased from ThermoFisher.

### Plasmids used

| Plasmid name | Addgene ID | Reference |
| --- | --- | --- |
| FLAG-S6K1 in pRK5 | 100509 | Gu et al. 2017 |
| FLAG-metap2 in pLJM60 | 100510 | Gu et al. 2017 |
| HA-metap2 in pRK5 | 100512 | Gu et al. 2017 |
| HA-GLUT5 in pRK5 |  | This study |
| FLAG-TKFC in pLJM60 |  | This study |
| FLAG-GK in pLJM60 |  | This study |
| GPI in pCW57.1 |  | This study |
| TPI in pCW57.1 |  | This study |
| FLAG(3x)-ALDOA in pCW57.1 |  | This study |
| FLAG(3x)-GAPDH in pCW57.1 |  | This study |
| HA-Mios in pRK5 |  | This study |
| WDR24 in pRK5 |  | This study |
| WDR59 in pRK5 |  | This study |
| SEC13 in pRK5 |  | This study |
| SEH1L in pRK5 |  | This study |

### Cell lines and tissue culture

HEK-293T were normally cultured in DMEM high glucose (25mM) with 10% IFS and supplemented with 2 mM glutamine. These cell lines were maintained at 37°C and 5% CO2. Cells were passaged every 48 hours and were not allowed to ever reach more than 90% confluency. For ALDO and GAPDH dox off experiments, cells were grown in DMEM low glucose (5mM).

### Transfections

For the transfection of cDNA expression constructs into HEK-293T cells, 1.5 – 2 million cells were seeded in 10 cm dishes. Using the polyethylenimine method (*39*), cells were transfected 24 hours after seeding with the indicated pRK5 based expression vectors. Experiments were done 36-48 hours after transfection. The total amount of DNA transfected was normalized to 5 μg with the empty pRK5 vector. The following amounts of cDNA were used in the indicated figures.

Fig1C: 2ng of FLAG-S6K (pRK5) + 500ng (HA-metap2 in pRK5 or HA-GLUT5 in pRK5) Supp_Fig 7F: 2ng of FLAG-S6K (pRK5) + either 2ug HA-Metap2 (pRK5) or 800ng each of HA-Mios, WDR24, WDR59, and 1200ng SEC13 and SEH1L.

### Lentiviral production and lentiviral infections

HEK-293T cells were seeded at a density of 750,000 cells per well of a 6-well plate in IMDM with 20% IFS. 24 hours after seeding, VSV-G envelope and CMV ΔVPR packaging plasmids were co-transfected with either pLJM60 containing cDNAs, pLentiCRISPRv2 with indicated guide sequences, or pCW57.1_tTA with the indicated cDNA, using XTremeGene 9 transfection reagent (Roche). 12 hours after transfection, the media was changed to fresh IMDM 20% IFS. 36 hours after the media change, the virus-containing supernatant was collected and passed through a 0.45 μm filter. Target cells were plated in 6-well plates with 8 μg/mL polybrene and incubated with virus-containing media. Cells were spinfected at 2200 rpm for 45 minutes at 37°C. 12 hours later, the virus containing media was replaced with fresh DMEM media. 24-48 hours after spinfection, cells were passaged into puromycin for pLJM60 or pLentiCRISPR or blasticidin for pCW57.1_tTA.

### Nutrient starvation experiments

Cells were seeded at 1 million cells per well in a 6-well plate format in fibronectin coated wells the day before the experiment. Cells were washed 1x with 1 mL of the starvation media (-glucose RPMI, -leucine RPMI, or - AA RPMI) and starved in 1 mL of the same starvation media or starved and restimulated with glucose or the indicated nutrient. For glyceraldehyde (GA) and dihydroxyacetone (DHA) restimulations, 500 mM solutions of GA and DHA in water were prepared immediately before adding it to cells.

### Cell lysis and immunoprecipitations

Cells were rinsed with cold PBS and lysed in lysis buffer (1% Triton, 10 mM β-glycerol phosphate, 10 mM pyrophosphate, 40 mM Hepes pH 7.4, 2.5 mM MgCl2 and 1 tablet of EDTA-free protease inhibitor [Roche] (per 25 ml of buffer)). Cell lysates were cleared by centrifugation in microcentrifuge (15,000 rpm for 10 minutes at 4°C). Cell lysate samples were prepared by addition of 5X sample buffer (0.242 M Tris, 10% SDS, 25% glycerol, 0.5 M DTT, and bromophenol blue), resolved by 8%-16% SDS-PAGE, and analyzed by immunoblotting.

For anti-FLAG immunoprecipitations, anti-FLAG M2 Affinity Gel (Sigma A2220) was washed with lysis buffer three times and then resuspended to a ratio of 50:50 affinity gel to lysis buffer. 25 μL of a well-mixed slurry was added to cleared lysates and incubated at 4°C with shaking for 90-120 minutes. Immunoprecipitates were then washed three times, once with lysis buffer and twice with lysis buffer with 500 mM NaCl. Immunoprecipitated proteins were denatured by addition of 50 μL of SDS-containing sample buffer (0.121 M Tris, 5% SDS, 12.5% glycerol, 0.25 M DTT, and bromophenol blue) and heated in boiling water for 5 minutes. Denatured samples were resolved by 8%-12% SDS-PAGE, and analyzed by immunoblotting.

### Generation of knock-out cell lines using CRISPR-Cas9 technology

To generate CRISPR/Cas9-mediated gene knockouts in HEK-293T cells, the following guide sequences were used to target each gene. For clonal KOs, guides were cloned into pX330 and for stable overexpression into pLentiCRISPR, as previously described.^34^

AAVS1: GGGGCCACTAGGGACAGGAT

AMPKα1 (PRKAA1): GAAGATCGGCCACTACATTC

AMPKα2 (PRKAA2): GAAGATCGGACACTACGTGC

FLCN: GGAAGGGCCAGGAGTTGATG

NPRL3: GCTGCACTCACCATCAGCCA

G6PD: GACACACTTACCAGATGGTG

Mios: ATCACATCAGTAAACATGAG

Axin1_sg1: GGAGCCTCAGAAGTTCGCGG

Axin1_sg2: GGAGCTCATCCACCGCCTGG

GPI: CAACCATGGGCATATCCTGG

ALDOA: CATTGGCACCGAGAACACCG

ALDOB: AAAACACTGAAGAGAACCGC

ALDOC: GGCTGGGTACGAGTGAGGCA

TPI: TGTCTTTGGGGAGTCAGATG

GAPDH: TGCTGGCGCTGAGTACGTCG

GPD1_sg1: AGAATGTCAAATACCTGCCA

GPD1_sg2: AATACCCACATGGTCACCCG

GPD1L: GAGAGTGCCCAAGAAAGCGC

GPD2_sg1: GGGACGATTCTTGTTGGAGG

GPD2_sg2: GATATCCTTGTTATTGGAGG

KPTN: ATCACATCAGTAAACATGAG

C12orf66: GGCTAAGGACAATGTGGAGA

On day one, 2 million HEK-293T cells were seeded in a 10-cm plate. Twenty-four hours after seeding, each well was transfected with 1 μg shGFP pLKO, 1 μg of the pX330 guide construct and 3 μg of empty pRK5 using XtremeGene9. Two days after transfection, cells were moved to a new 10-cm plate into puromycin containing media. Forty-eight hours after selection, the media was switched to media not containing puromycin. Cells were allowed to recover for 1 week after selection prior to single-cell sorting with a flow cytometer into the wells of a 96-well plate containing 150 μl of DMEM supplemented with 30% IFS.

### Generation of Dox-off cell lines

cDNAs for human GPI, ALDOA, TPI, or GAPDH were cloned from HEK-293T cDNA. The cDNAs were made resistant for their respective sgRNA as follows. GPI: sgRNA sequence <CAACCATGGGCATATCCTGG> was used; the PAM sequence in the cDNA was mutated by mutating the codon for valine 53 from GTG>GTA using overhang extension PCR. ALODA: sgRNA sequence < CATTGGCACCGAGAACACCG> was used; the PAM sequence in the cDNA was removed by mutating the codon for glutamate 72 from GAG>GAA. TPI: Based on molecular weight, HEK-293T cells express isoform 2 (Uniprot: P60174-1) of TPI1. The sgRNA < TGTCTTTGGGGAGTCAGATG> was used, the cDNA was naturally resistant to this guide because the PAM sequence is partially in an intron, therefore no modification was made for this cDNA. GAPDH: sgRNA sequence < TGCTGGCGCTGAGTACGTCG> was used; the PAM sequence in the cDNA was removed by mutating the codon for valine 96 from GTG>GTA.

Cells were transduced with lentivirus produced from the pCW57 vector encoding the dox-off cDNA and a blasticidin resistance gene and were selected for 7 days with blasticidin. Next, cells were then either transduced to express stable Cas9 and the respective sgRNA (GPI, TPI, and GAPDH) or for ALDOA guides against ALDOA, ALDOB, and ALDOC were co-transfected along with an shGFP pLKO vector encoding a puromycin resistance gene. Cells were puromycin selected and allowed to expand for an additional least 7 days. Cells were single cell sorted and screened by western blotting for the appropriate dox-off status.

### Doxycyline Treatment

Doxycyline was prepared as a 30 ug/mL stock solution and aliquots were stored at −80°C. Aliquots are good for 2-3 months at −80°C. Cells were treated with 30 ng/mL doxycycline and cultured using 5mM glucose DMEM for 5 days for the GAPDH and ALDOA experiments. For GPI and TPI, cells were treated for at least 9 days prior to the experiment.

Notably, when >3 mM glucose was added to cells lacking GAPDH expression, it caused an 80% decrease in ATP levels 3 mM glucose and 30-fold increase in AMP levels. This is likely due to the fact that glucose is phosphorylated twice in upper glycolysis, but the phosphates are not liberated as this occurs after the GAPDH step of glycolysis. Therefore, when GAPDH is absent, glucose acts as an ATP-sink. Thus, it is hard to interpret the signaling status of cells in the presence of >3 mM glucose and lacking GAPDH expression because large decreases in ATP levels are known to decrease mTORC1 activity due to its relatively high Km for ATP of 1mM^35^.

### Koningic Acid Treatment

Koningic acid was ordered in a 250 ug size and resuspended in 178 uL of sterile water to make a 5 mM stock solution. The stock solution can be aliquoted and stored at −80° C for 2 weeks at most. Cells were starved of glucose and KA was added as indicated. KA if added to glucose replete cells was very toxic so it was only added to glucose-starved cells where it was well tolerated during the 3-hour incubation period. Similar to cells lacking GAPDH expression, glucose addition to KA treated cells caused a marked decrease in ATP levels and decrease in mTORC1 signaling. This effect is likely similar to the phenomenon observed in cells lacking GAPDH expression and to that observed by Dennis et al^35^.

### Immunofluorescence assays

For the experiment in Figure 4E, 400,000 HEK-293T cells were plated on fibronectin-coated glass coverslips in 6-well tissue culture plates. After 24 hours, the slides were rinsed once with PBS and fixed with 4% paraformaldehyde in PBS for 15 minutes at room temperature. The slides were then rinsed three times with PBS and the cells permeabilized with 0.05% Triton X-100 in PBS for 5 minutes at room temperature. The slides were rinsed three times with PBS and then blocked for 1 hour in Odyssey blocking buffer at room temperature. The slides were incubated with primary antibody in Odyssey blocking buffer at 4°C overnight, rinsed three times with PBS, and incubated with secondary antibodies produced in donkey (diluted 1:1000 in Odyssey blocking buffer) for 50 minutes at room temperature in the dark, and washed three times with PBS. The primary antibodies used were directed against mTOR (CST; 1:100-1:300 dilution), LAMP2 (Santa Cruz Biotechnology; 1:300 dilution). Slides were mounted on glass coverslips using Vectashield (Vector Laboratories) containing DAPI.

Images were acquired on a Zeiss AxioVert200M microscope with a 63X oil immersion objective and a Yokogawa CSU-22 spinning disk confocal head with a Borealis modification (Spectral Applied Research/Andor) and a Hamamatsu ORCA-ER CCD camera. The MetaMorph software package (Molecular Devices) was used to control the hardware and image acquisition. The excitation lasers used to capture the images were 488 nm (LAMP2) and 561 nm (mTOR).

Lysosomal enrichment was quantified as previously described^10^. Raw image files were opened in the Fiji software and a maximum-intensity projection of a Z stack of ~6–8 contiguous focal planes (∼0.5μm each) was used. In each cell analyzed, a cytoplasmic region of interest containing lysosomes was chosen by finding a punctum of high LAMP2 signal and in this area the mean fluorescence intensities (MFIs) of the 488 nm (LAMP2) and 561 nm channels (mTOR) were measured. In the same cell an equivalently sized area in a region of the cytoplasm not containing lysosomes with low LAMP2 signal was chosen and the MFIs of the 488nm and 561nm channels were also measured in this area. For each channel, the MFI of the non-lysosomal area was subtracted from that of the lysosomal area. The value obtained for the 561nm channel was then divided by the analogous value for the 488nm channel to give the lysosomal enrichment factor shown in the bar graphs in the figures. A lysosomal enrichment factor close to 1 indicates that the mTOR (561 nm) signal was enriched in a region of the cell containing lysosomes over one that does not. A lysosomal enrichment factor closer to 0 indicates that the mTOR(561 nm) signal was not enriched at the lysosomes, indicating no specific co-localization with the LAMP2 (488 nm).

### Extraction of metabolites for LC/MS analyses

Cells were seeded in fibronectin coated 6 well plates at a density of 1 million cells per well the day before the experiment. On the day of the experiment, cells were starved or starved and restimulated as indicated. At the time of lysis, the media was aspirated, cells were washed once with 1mL of cold saline, and metabolites were extracted by adding 800 uL of cold 80% methanol containing 500 nM isotope-labeled internal standards. Methanol extracts were moved to pre-chilled Eppendorf tubes and samples were moved immediately to dry ice. Samples were briefly vortexed for 1 min, spun 15,000 rpm at 4°C for 10 minutes. The supernatant was dried by vacuum centrifugation and stored at −80°C. Just before LC/MS analysis, samples were resuspended in 50uL of LC/MS grade water, cleared of any insoluble debris by centrifugation at 15,000rpm.

### HILIC LC/MS

Polar metabolite profiling was conducted on a QExactive bench top orbitrap mass spectrometer equipped with an Ion Max source and a HESI II probe, which was coupled to a Dionex UltiMate 3000 HPLC system (Thermo Fisher Scientific, San Jose, CA). External mass calibration was performed using the standard calibration mixture every 7 days. Typically, samples were reconstituted in 100 uL water and 2 uL were injected onto a SeQuant® ZIC®-pHILIC 150 × 2.1 mm analytical column equipped with a 2.1 × 20 mm guard column (both 5 mm particle size; EMD Millipore). Buffer A was 20 mM ammonium carbonate, 0.1% ammonium hydroxide; Buffer B was acetonitrile. The column oven and autosampler tray were held at 25°C and 4°C, respectively. The chromatographic gradient was run at a flow rate of 0.150 mL/min as follows: 0-20 min: linear gradient from 80-20% B; 20-20.5 min: linear gradient form 20-80% B; 20.5-28 min: hold at 80% B. The mass spectrometer was operated in full-scan, polarity-switching mode, with the spray voltage set to 3.0 kV, the heated capillary held at 275°C, and the HESI probe held at 350°C. The sheath gas flow was set to 40 units, the auxiliary gas flow was set to 15 units, and the sweep gas flow was set to 1 unit. MS data acquisition was performed in a range of *m/z* = 70–1000, with the resolution set at 70,000, the AGC target at 1×106, and the maximum injection time (Max IT) at 20 msec. To increase sensitivity to DHAP/GAP, a targeted selected ion monitoring (tSIM) scan in negative mode was included. The isolation window was set at 1.0 *m/z* and tSIM scans were centered at *m/z* = 168.99080. Relative quantitation of polar metabolites was performed with XCalibur QuanBrowser 2.2 and TraceFinder 4.1 (both Thermo Fisher Scientific) using a 5 ppm mass tolerance and referencing an in-house library of chemical standards.

### Aniline labeling of metabolite extracts for metabolomics

We adapted a previously published method in order to measure aniline-GAP (glyceraldehyde 3-phosphate) adducts^36,37^. Following methanol extraction, 300 uL of the ~800 uL samples were used in the aniline reaction. Both aniline and EDC solutions were prepared fresh in water. For the aniline solution, 777.5 mg/mL solution (6 M) was prepared in water and 6 uL of 10 M NaOH was added per 100 uL of the solution to increase pH to 4.5. To 300 uL of methanol extract, 30 uL of 6M Aniline-HCl and 30 uL of 200 mg/mL EDC solution were added. Samples were gently vortexed for 2 hours at room temperature. At the end of the two hours 5 uL of triethylamine was added to stop the reaction. Samples were then dried by vacuum centrifugation and stored at −80°C. Just before LC/MS analysis, samples were resuspended in 50 uL of LC/MS grade water, cleared of any insoluble debris by centrifugation at 15,000 rpm.

### C8 LC/MS

The LC and mass spectrometer general settings were as described above. Typically, 10 uL of a sample was injected onto a Kinetex C8 2.6 μm, 2.1 × 30 mm column (Phenomenex). Mobile Phase A was 0.1% formic acid in water and Mobile Phase B was 0.1% formic acid in acetonitrile. The column oven was set to 25°C and the flow rate was 0.250 mL/min. The gradient was as follows: 0-1 min: hold at 5% B; 1-6 min: linear gradient 5-70% B; 6-8 min: linear gradient 70-100% B; 8.1-10 min: hold at 5% B. The data were acquired in negative ion mode with a scan range of *m/z* = 140-380. A tSIM was included for the doubly labeled GAP-aniline adduct and was centered on *m/z* = 319.08420. Relative quantification of GAP-aniline was performed with XCalibur QuanBrowser 2.2 and TraceFinder 4.1 (both Thermo Fisher Scientific) using a 5 ppm mass tolerance.

### Statistical analyses

Two-tailed t tests were used for comparison between two groups. Generally sample n = 3 representing three biological replicates. Reported p-values were not adjusted for multiple hypothesis testing. Therefore, some comparisons deemed to be statistically significant with p-values <0.05 may reflect type I error.

## Acknowledgements

The authors would like to thank Max Valenstein, Jessica Spinelli, and all the current members of the Sabatini lab for helpful insights. This work was supported by grants from the NIH (R01 CA103866, R01 CA129105, and R37 AI047389). J.M.O. was supported by a fellowship grant F30CA210373 from the National Cancer Institute and the Harvard-MIT MSTP training grant T32GM007753 from the National Institute of General Medical Sciences. P.A.K. was supported by a scholarship from Santander Universidades Mobility Fund granted by Adam Mickiewicz University in Poznan, Poland. A.L.C. was supported by a fellowship grant F31DK113665 from the National Institute of Diabetes and Digestive and Kidney Diseases. D.M.S. is an investigator of the Howard Hughes Medical Institute and an American Cancer Society Research Professor.

## Author contributions

J.M.O. and D.M.S. initiated the project and designed the research plan with input from P.A.K. J.M.O. and P.A.K. conducted experiments and analyzed data with technical assistance from SMS. S.H.C., T.K., and C.A.L. operated the LC/MS platform and assisted in the method development to measure GAP-aniline. J.M.O. and D.M.S. wrote the manuscript with assistance from P.A.K. and C.A.L.

## Conflicts of interest

DMS is a founder, shareholder, and a member of the scientific advisory board for Navitor Pharmaceuticals, which targets the mTORC1 pathway for therapeutic benefit. J.M.O. is a shareholder of Navitor Pharmaceuticals.

## Notes

### Competing Interest Statement

DMS is a founder, shareholder, and a member of the scientific advisory board for Navitor Pharmaceuticals, which is targeting the mTORC1 pathway for therapeutic benefit. J.M.O. is a shareholder of Navitor Pharmaceuticals. The other authors declare no competing intersts.

